# Cognitive and brain development is independently influenced by socioeconomic status and polygenic scores for educational attainment

**DOI:** 10.1101/866624

**Authors:** Nicholas Judd, Bruno Sauce, John Wiedenhoeft, Jeshua Tromp, Bader Chaarani, Alexander Schliep, Betteke van Noort, Jani Penttilä, Yvonne Grimmer, Corinna Insensee, Andreas Becker, Tobias Banaschewski, Arun L.W. Bokde, Erin Burke Quinlan, Sylvane Desrivières, Herta Flor, Antoine Grigis, Penny Gowland, Andreas Heinz, Bernd Ittermann, Jean-Luc Martinot, Marie-Laure Paillère Martinot, Eric Artiges, Frauke Nees, Dimitri Papadopoulos Orfanos, Tomáš Paus, Luise Poustka, Sarah Hohmann, Sabina Millenet, Juliane H. Fröhner, Michael N. Smolka, Henrik Walter, Robert Whelan, Gunter Schumann, Hugh Garavan, Torkel Klingberg

## Abstract

Genetic factors and socioeconomic (SES) inequalities play a large role in educational attainment, and both have been associated with variations in brain structure and cognition. However, genetics and SES are correlated, and no prior study has assessed their neural associations independently. Here we used polygenic score for educational attainment (EduYears-PGS) as well as SES, in a longitudinal study of 551 adolescents, to tease apart genetic and environmental associations with brain development and cognition. Subjects received a structural MRI scan at ages 14 and 19. At both time-points, they performed three working memory (WM) tasks. SES and EduYears-PGS were correlated (r = 0.27) and had both common and independent associations with brain structure and cognition. Specifically, lower SES was related to less total cortical surface area and lower WM. EduYears-PGS was also related to total cortical surface area, but in addition had a regional association with surface area in the right parietal lobe, a region related to non-verbal cognitive functions, including mathematics, spatial cognition, and WM. SES, but not EduYears-PGS, was related to a change in total cortical surface area from age 14 to 19. This is the first study demonstrating a regional association of EduYears-PGS and the independent prediction of SES on cognitive function and brain development. It suggests that the SES inequalities, in particular parental education, are related to global aspects of cortical development, and exert a persistent influence on brain development during adolescence.

**Significance statement:** The influence of socioeconomic (SES) inequalities on brain and cognitive development is a hotly debated topic. However, previous studies have not considered that genetic factors overlap with SES. Here we showed, for the first time, that SES and EduYears-PGS (a score from thousands of genetic markers for educational attainment) have independent associations with both cognition and global cortical surface area in adolescents. EduYears-PGS also had a localized association in the brain: the intraparietal sulcus, a region related to non-verbal intelligence. In contrast, SES had global, but not regional, associations, and these persisted throughout adolescence. This suggests that the influence of SES inequalities is widespread – a result that opposes the current paradigm and can help inform policies in education.

## Introduction

Adolescence is a critical phase in neural and cognitive development during which much of adult trajectories are shaped, yet there is still much we don’t know about the environmental and genetic influences during this period ^1–3^.

Socioeconomic status (SES) inequalities have been associated with differences in executive function, memory, emotional regulation and educational attainment ^1–3^. SES is also associated with functional and structural neural differences in a wide range of cortical areas, including those underlying higher cognitive functions ^4–8^. Although SES is commonly assumed to represent a purely environmental factor, large portions of variability in SES can be explained by additive genetic factors ^9^. SES explains 9-25% of educational achievement, half of which is suggested to be genetically mediated ^10,11^. Genetic differences are thus a confound not adequately addressed by prior studies of SES and neural development ^12^.

Recent advances in behavioral genetics have identified substantial genetic associations with educational attainment. In a genome-wide association study with 1.1 million individuals, Lee and colleagues ^13^ described a polygenic score (EduYears-PGS) which explained up to 13% of the variance in educational attainment. That represents the best composite of genetic variants currently known for educational attainment, and many of these variants are relevant for brain development and show tissue-specific expression in the cerebral cortex. Therefore, while SES is a strong environmental predictor for educational attainment, EduYears-PGS is a powerful genetic predictor. And at least some of the impact from these two predictors is likely to overlap and to be via cognition.

Here we evaluated the independent associations of SES and EduYears-PGS in a longitudinal study of brain development and cognitive function in 551 adolescents. A major strength of our sample is that it was specifically designed to include poor communities for an accurate range of SES inequalities ^14^. Adolescent age was around 14 at the first timepoint, and 19 at the second time point, a time-span that captures secondary education. We chose working memory (WM) as a measure of cognitive function since it is highly correlated with academic ability and was available at both timepoints ^15^.

Genetics and environment effects could either be manifested in local cortical regions, such as a specific prefrontal area, or they could have global effects, for example via a molecular mechanism that involves most cortical neurons. The distinction is important because it tells us about the potential mechanisms in play, has functional consequences for the individual and could have implications for remedial interventions. Therefore, we tested for distinct global and regional associations – a practice still ignored in the literature ^4,5,8^, with the exception of only one known study to date ^16^. The latter, however, did not control for any genetic component of SES.

We defined SES as a combination of income, education, and neighborhood quality. A single, combined SES component allowed for better comparison with EduYears-PGS (itself a single, genetic composite) and was hypothesized to increase power to detect associations in case of additive influences from each SES-related factor. A combined measure also makes our results more interpretable in light of the broader SES literature (most of it also uses composites scores). Yet, recent research has highlighted issues and limitations arising from using SES as a composite measure ^17^. Therefore, in a post hoc analysis, we split our SES composite into parental education, income-related stresses, and neighborhood quality to evaluate how these components contributed to our main findings from the analysis of the combined SES measure.

We used a bivariate latent change score model (bLCS) to analyze the independent relationships of EduYears-PGS and SES on cognition and global measures of cortical thickness and surface area at age 14 and on the amount of change until 19 while controlling for age, gender, and scan site. Second, to examine regional associations in the cortex, we used cluster corrected vertex-wise analyses to isolate independent variation related to SES and EduYears-PGS, while controlling for respective global values.

## Results

### Cognitive associations in Early Adolescence

Data from 551 adolescents recruited by the IMAGEN consortium (https://imagen-europe.com) were included. First, global measures of cortical surface area and scores from three WM tasks were entered into a bLCS model.

A strict measurement invariant, bivariate-LCS model (Fig. 1), with SES and EduYears-PGS as covariates of interest, fit the data well, RMSEA = .031, CFI = .997. We found that SES and EduYears-PGS were significantly and positively correlated (r = .27, p < .001). SES had a positive and significant association to WM, even when correcting for EduYears-PGS (β = .23, p < .001). Splitting SES into subcomponents revealed that only parental education was significantly associated with WM. EduYears-PGS also had a positive, independent, although weaker relationship, with WM at 14 (β = .11, p < .05).

**Figure 1.**
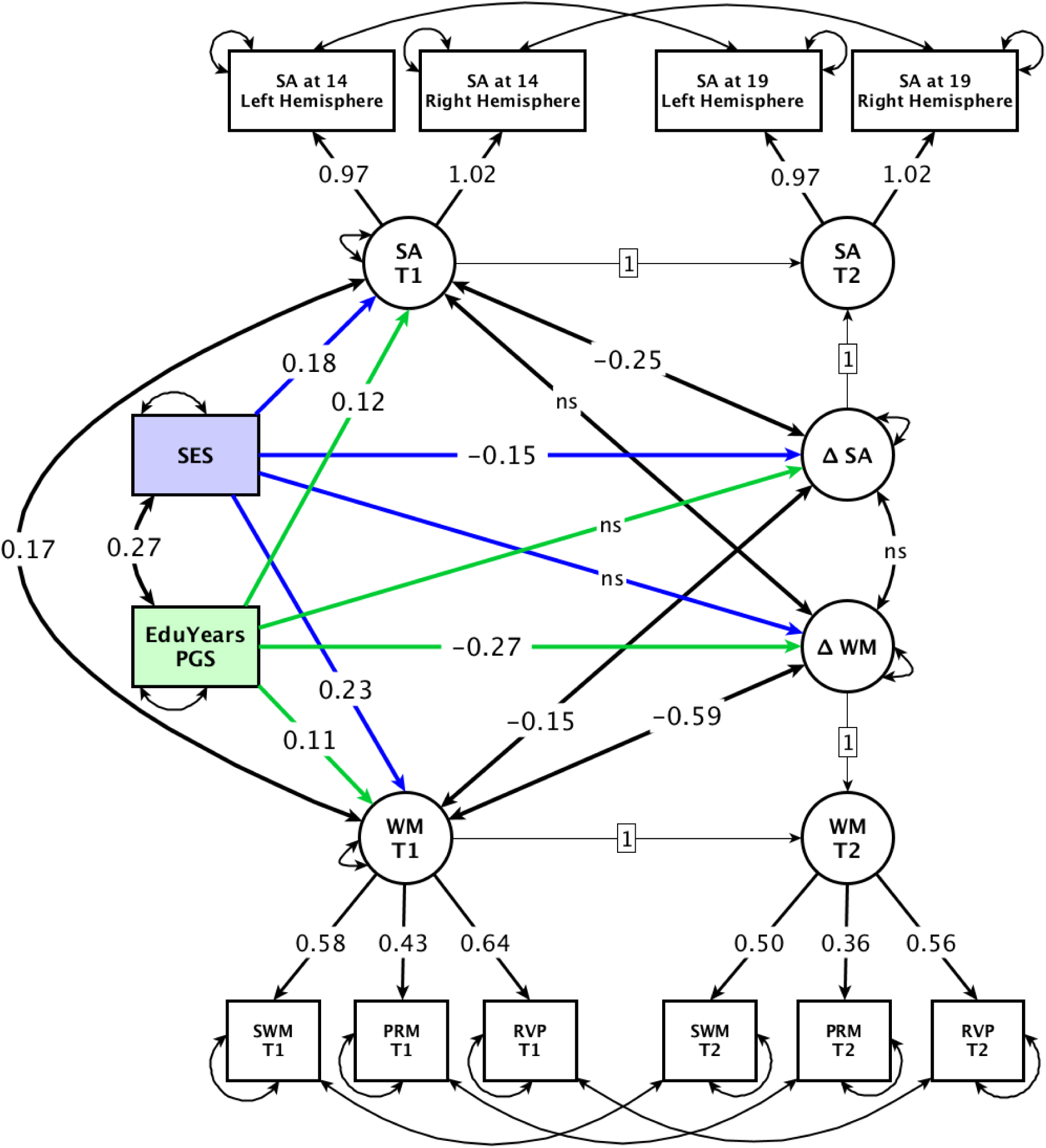
Path diagram of a strict measurement invariant bLCS model with the change of surface area (SA) and working memory (WM) from 14 to 19. SES and EduYears-PGS are our exogenous variables of interest, all variables are standardized. Following convention, squares represent observed variables and circles represent latent variables. Single headed arrows denote regressions while double headed arrows represent variances, co-variances or error. See SI Appendix, Fig. S2 for a specification without SES and EduYears-PGS and with SA and WM at 14 as exogenous variables on change.

Two subtests of an IQ test (WISC, perceptual reasoning and verbal comprehension) were available when participants were 14 years old, but not at 19. The results from both subtests mirrored the results from WM with significant, independent relationships for both SES and EduYears-PGS (SI Appendix, Tables S1 and S2).

### Global Surface Area in Early Adolescence

Global surface area in 14-year-olds was significantly associated with both SES (β = .18, p < .001) and EduYears-PGS (β = .12, p < .01), each contributing unique variance. Both associations were positive with a stronger association for SES (Fig. 2). As expected, surface area and WM were both correlated at age 14 (r = .17, p < .01). A linear model showed a relationship between surface area and gender, yet there was no interaction between gender and SES (see SI Appendix, Table S3).

**Figure 2.**
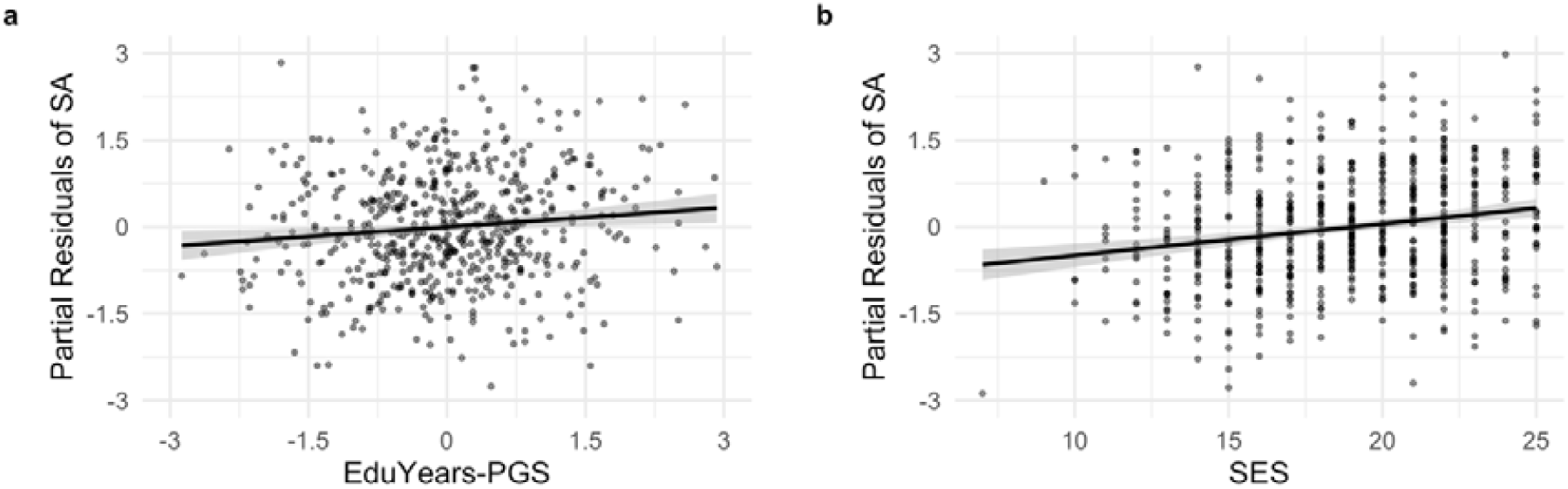
Partial residual plots of a) EduYears-PGS and b) SES on global surface area (SA) at age 14.

An important question on SES and brain development is whether results only hold for certain SES levels or whether it is linear for all ranges of SES. To evaluate this, we fitted models with the natural logarithmic and a 3^rd^ degree polynomial for SES. In comparison to the linear model (AIC = −1161), neither the logarithmic (AIC = −1161) nor the polynomial model (AIC = −1157) showed improved model fits. There was thus no evidence of non-linearity in our sample.

### Regional cortical associations in Early Adolescence

Next, we investigated regional cortical associations of SES and EduYears-PGS with surface area at age 14. In this analysis, we corrected for total surface area, age, sex and scanning site. A vertex-wise analysis (CFT < .001, CWP < .05) identified one cluster, uniquely related to EduYears-PGS, located in the right intraparietal sulcus, partly covering the top of the supramarginal gyrus (309 mm^2^, [x = 49, y = −42, z = 39], p < .05) (Fig. 3).

**Figure 3.**
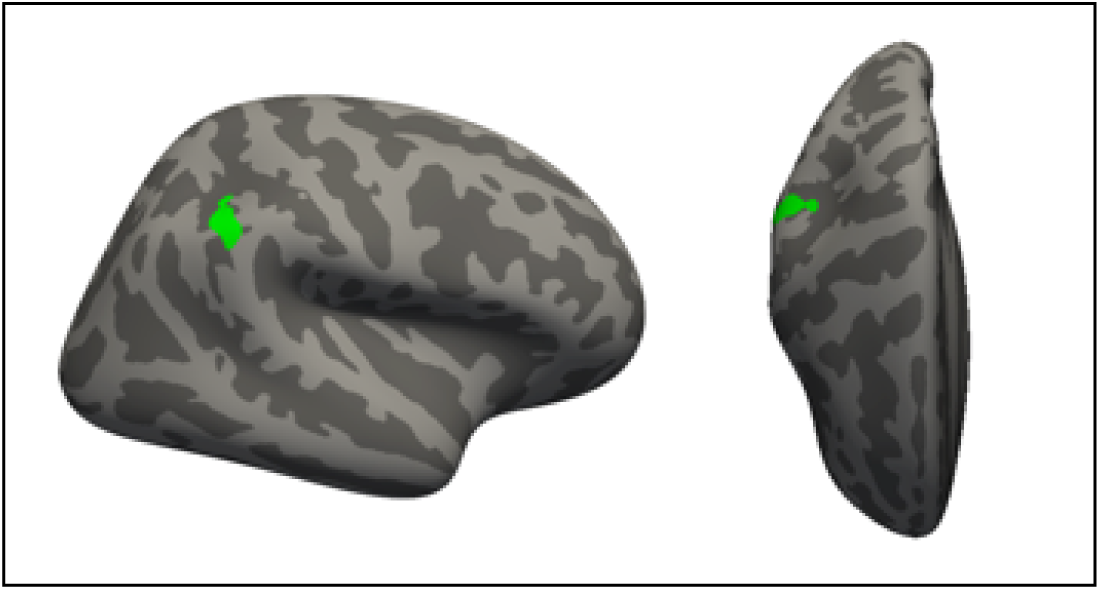
The right IPS in which surface area at 14 relates to EduYears-PGS while controlling for SES and global effects.

SES did not have any regional associations at age 14 when covarying for total surface area. Figure 4 illustrates this finding by showing uncorrected, vertex-wise, surface maps in which the mean image of all subjects 1 SD above the means for SES and EduYears-PGS (analyses are done separately for the two measures) are subtracted from the mean image of all subjects 1 SD below the mean (SES-low n=83, SES-high n=86, PGS-low n=79, PGS-high n=80). To obtain representative mean and standard deviations of random data, this procedure was repeated 10,000 times on randomly sampled data without replacement (groups n = 85). Consistent with the statistical analyses, these maps and histograms suggest that almost all the cortex was related to SES inequalities. Firstly, this finding is clearly shown by the density histogram of SES, in which almost every vertex value is above zero (i.e., the mean of randomly sampled data) (Fig. 4a). To further illustrate this widespread cortical association, we took all vertex values from one hemisphere to produce a square, 2-dimensional flat-map which contained all data from the hemisphere in one matrix (Fig. 4b). We thereby reduced topographical information (i.e., localization to brain regions) in order to only focus on size and distribution of relative increases and decreases in surface areas. This resulted in 3 flat maps (random, EduYears-PGS and SES), the color bar was constrained to the minimum and maximum values of random data.

**Figure 4.**
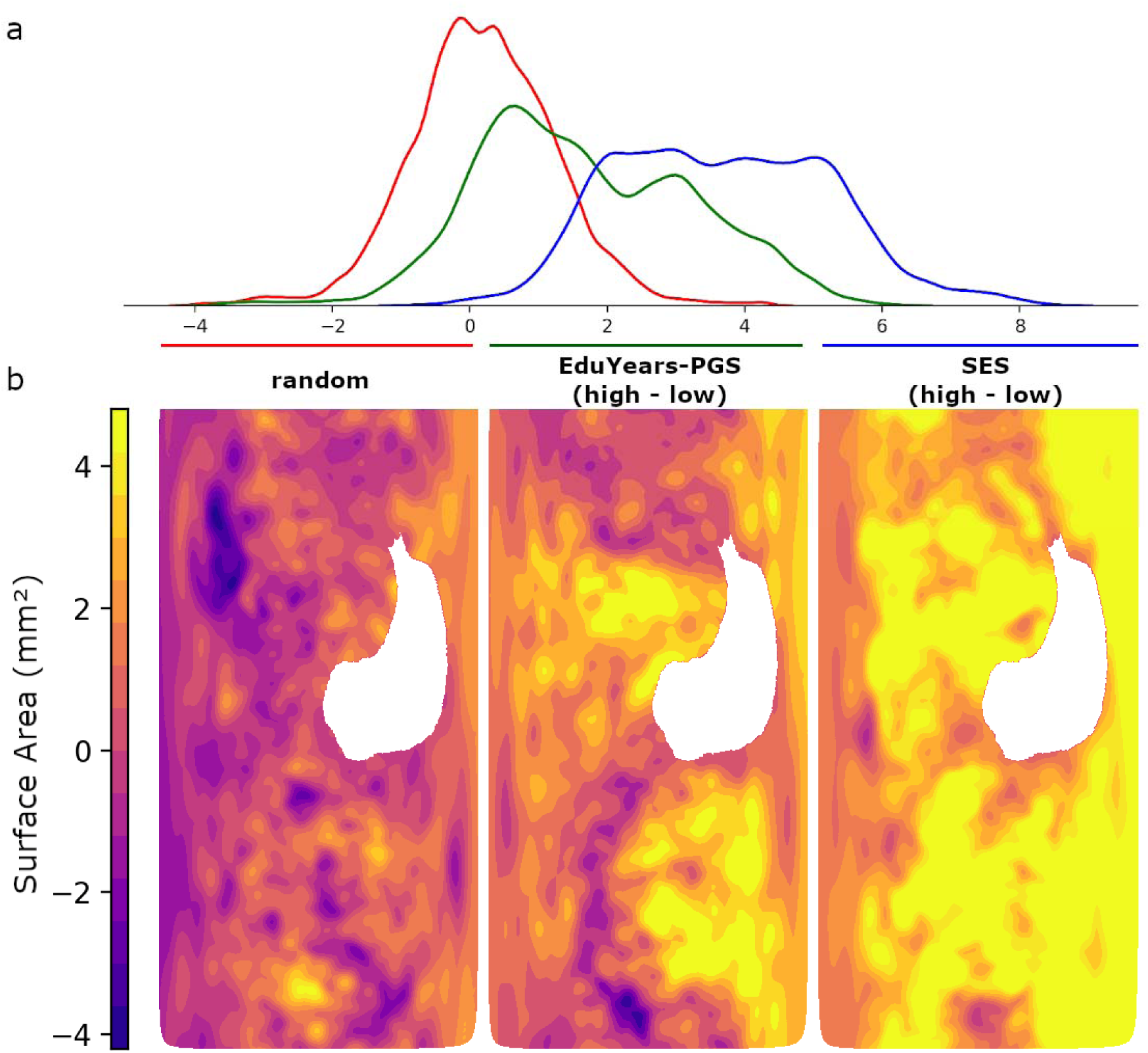
Global surface area of the left hemisphere displayed with **(a)** density histogram and, (**b**) flat maps. White area corresponds to missing values from the corpus callosum. Data for EduYears-PGS (middle figure) and SES (right most) were calculated by vertex-wise averaging of subjects 1 SD above the mean subtracted from those 1 SD below. Random data were selected by random sampling without replacement (n = 85 for each group). The two groups were then separately averaged vertex-wise and subtracted from each other. This was repeated 10,000 times to calculate an average random mean and standard deviation. To produce plots for random data the first representative sample was chosen.

### Adolescent Development in Cognition

There was significant improvement in WM from age 14 to 19 (β = 2.31, p < .001) and this was more pronounced in the subjects with lower WM scores at age 14 (r = −0.59, p < .01). However, inspection of the distributions and variation of performance on the WM tasks at the second timepoint suggested that there were ceiling effects, which could artificially lead to subjects who performed well at age 14 having less room for improvement. SES was not related to change in WM (p > .05), but EduYears-PGS was (β = -.27, p < .05).

### Adolescent Development in Surface Area

Next, we investigated global changes in surface area over the course of five years. Correlation in global surface area between the ages of 14 (Mean = 181682mm^2^, SD = 16130mm^2^) and 19 (Mean = 177625mm^2^, SD = 15860mm^2^) was very high (r = .99, p < .001). On average, there was a significant decrease in total surface area over five years (−4057mm^2^, p < .001, paired t-test; SI Appendix, Fig. S6a). SES, but not EduYears-PGS, was significantly related to change in surface area (β = -.15, p < .01).

Subtracting each individual’s surface area at age 14 from their surface area at age 19 removes inter-individual differences in total surface area. However, prior studies have shown that change is dependent on point of departure, as subjects with higher initial surface area show larger change ^18^. Therefore, we performed regional analyses both with and without correction for total surface area at 14. Using vertex-wise subtracted images, we found a cluster in left caudal superior frontal sulcus (138 mm^2^, [x = −28, y = 10, z = 48], p < .05) significantly associated with SES (SI Appendix, Fig. S4). This cluster was no longer significant when covarying for global surface area at 14. Furthermore, in a post hoc analysis we found no significant association of change in this region to the change of WM. No clusters were found for the relationship of EduYears-PGS on change in cortical surface area.

### Cortical thickness

Global cortical thickness decreased from age 14 to 19 (−.12mm, p < .001, paired t-test; SI Appendix, Fig. S6b). A strict measurement invariant, bivariate-LCS model for global cortical thickness (SI Appendix, Fig. S1b), with SES and EduYears-PGS as covariates of interest, fit the data well, RMSEA = .037, CFI = .988. Neither SES nor EduYears-PGS were significantly related to average cortical thickness at age 14 or to the thinning over adolescence. Furthermore, there were no significant regional associations (at 14 or from 14-19) for cortical thickness from either SES or EduYears-PGS, regardless of global correction.

### Amygdala and Hippocampal Volume

The main focus of this study was on neocortical development, with a specific focus on distinguishing between cortical thickness and cortical surface area. Although our method did not allow us to separate thickness and area for the amygdala and hippocampus, we analyzed the volume of these structures to tie into the wider body of SES literature.

The volume of the amygdala did not change from age 14 to 19 (13.6 mm^3^, p > .05, paired t-test; SI Appendix, Fig. S6c). A bivariate-LCS model for the amygdala with SES and EduYears-PGS, fit the data well, RMSEA = .025, CFI = .994. Neither SES nor EduYears-PGS were significantly related to amygdala volume at age 14 or to the change in volume over adolescence.

The hippocampus also did not show a significant change in volume over adolescence (−9.3mm^3^, p > .05, paired t-test; SI Appendix, Fig. S6d). A bivariate-LCS model (RMSEA = .035, CFI = .991) found hippocampal volume at age 14 to be related to SES (β = .12, p < .01), yet not EduYears-PGS. Hippocampal volume change over adolescence was not related to SES or EduYears-PGS.

### SES subcomponent analysis

As described in the introduction, our main analysis used an SES composite, which has advantages and limitations. Therefore, we post hoc split our SES composite into parental education, income-related stresses, and neighborhood quality. Each SES subcomponent was added individually in bivariate-LCS models with surface area, cortical thickness, amygdala volume and hippocampal volume. All models fit the data well (RMSEA < .08 and CFI > .95). All cortical and cognitive results that were significant for the SES composite were also significant when only using parental education as a covariate, while income and neighborhood were not significant (see SI Appendix, Table S5). The only discrepancy between the findings for parental education and the SES composite was regarding the relationship of parental education on hippocampal volume at age 14, which did not survive multiple comparisons correction.

## Discussion

Here we showed that both environmental (SES) and genetic (EduYears-PGS) factors, each of which influence educational attainment, play an important role in cognitive and brain development during adolescence. This is the first study to test for and confirm independent, non-overlapping associations for SES and EduYears-PGS. We found that although genetic and environmental determinants of educational attainment are correlated, they carry independent influences on cognition and brain development. Both SES and EduYears-PGS were related to total cortical surface area at age 14, with SES having only a global association, while EduYears-PGS also had a regional association with cortical surface area in the right intraparietal sulcus. In analyzing developmental changes, we found that SES, but not EduYears-PGS, continued to be relevant for surface area change from 14 to 19 years.

### Cognition in early adolescence

Both SES and EduYears-PGS independently correlated with WM in early adolescence, with SES having about twice as strong of an influence. This independent SES result is similar to one found by a meta-analysis for educational achievement (r = .3) ^11^. The IQ-subtests displayed the same pattern, showing that the associations were not specific for WM, but likely reflect a general effect on cognition. This is consistent with the well-known correlation between educational attainment and IQ^19,20^ and underscore the value of WM as a suitable and meaningful measure of adolescent cognition and cognitive development.

### Brain structure in early adolescence

The strong relationship observed between SES and global surface area at age 14 is consistent with prior findings ^4,8^. There are multiple potential mechanisms mediating an effect of SES on brain development, including stress and glucocorticoids during pregnancy, toxins, premature delivery, maternal care, lack of cognitive stimulation and chronic stress during childhood and adolescence ^6,21^. Those factors might exert their influence via brain functioning and possible epigenetic mechanisms. Reward signals and selected epigenetic markers have indeed been discussed as possible malleable neurobiological markers being associated with cognitive capacity in adolescents ^22^. However, to date, one epigenome-wide analysis for educational attainment casts doubt on possible epigenetic markers for educational attainment ^23^. The global result we observed could come from one or several factors with a global impact or be the result of several regional effects that together impact most of the cortex with wide-ranging behavioral outcomes^24^.

After correcting for global surface area, there were no regional associations with SES. Similarly, no clusters were significant when total intracranial volume was corrected for instead of global surface area. Although several studies report regional findings of SES on surface area and cortical thickness ^4,8^, these studies did not correct for global associations. A post-hoc analysis of our data showed that if the global differences are not controlled for, several clusters reach significance, in agreement with prior studies (SI Appendix, Fig. S5).

Contrasting the surface area in high vs low SES subjects (Fig. 4), it is clear that almost the entire cortex is related to SES inequalities. Without total surface area correction the regional associations can give a false sense of localization ^25,26^. We view it as more accurate to describe the association of SES on surface area as global in nature, where regional effects over-and-above this global effect cannot be statistically distinguished from noise.

EduYears-PGS was associated with global surface area, consistent with prior findings showing a relationship between EduYears-PGS and intracranial volume ^27,28^. This was expected for our polygenic score for educational attainment, given that both intracranial volume and total surface area were also shown in the past to correlate with IQ ^18,29^. In addition, here we found that EduYears-PGS was related to regional surface area in the intraparietal sulcus.

Regional cortical findings are consistent with the fact that some of the genetic markers from the EduYears-PGS are associated with regional gene expression ^13^. The intraparietal sulcus is a region typically associated with nonverbal reasoning, visuospatial WM, and mathematics ^30–32^. Gray matter volume ^33,34^ and brain activity ^35–37^ of the intraparietal sulcus predicts current and future mathematical skills in children and adolescents.

A study spanning 6-18 year-olds found that the right anterior intraparietal sulcus was associated with visuo-spatial WM at younger ages, and later during development associated with mathematics, suggesting a functional plasticity of this region ^38^. Given that both non-verbal reasoning, WM and mathematics are predictors of future educational attainment, it is of particular note that our data show the intraparietal sulcus to be specifically associated with EduYears-PGS above and beyond the polygenic influences on global cortical surface area.

We found no association between EduYears-PGS or SES with cortical thickness, in agreement with some previous studies on SES ^4^, but not others ^5,8^. It is important to emphasize that our quality control procedure was very strict and previous literature has shown cortical thickness results to change based upon strictness ^39–41^. This reasoning, combined with our large sample size and our findings for surface area, lead us to interpret this as an important null result.

### Adolescent Development

Over the course of five years, we found a global decrease in surface area. The amount of decrease was related to SES, but not EduYears-PGS, showing a continuing relationship of SES on brain development during adolescence. Although SES’s association with surface area at age 14 was positive, this relationship flipped (i.e. became negative) for the change over five years (Fig. 1). This is likely related to the non-linear developmental trajectories of surface area during childhood and adolescence with an inverted u-shape (typically a loss of surface area starting in adolescence) where height and delay of the peak can differ between individuals as well as between brain region ^24,42,43^.

There was an association between SES and regional change in the left caudal middle frontal gyrus, but this did not survive correction for global surface area at age 14. The interpretation of this regional finding is therefore unclear.

One of the benefits of using a bLCS model is that it allows us to examine the influence of baseline measures (e.g., surface area and WM at 14) on the change of those measures.

Interestingly, higher WM, independent of surface area at 14, and after correction for SES and EduYears-PGS, was related to a decrease in global surface area during adolescence. Recent research has shown intra-individual change in cognition associated with later change in surface area ^44^. An accelerated reduction of surface area during development has previously been observed in higher IQ subjects ^18^. In summary, SA typically decreases during adolescence. Higher WM enhances this decrease, suggesting that it is a beneficial developmental process blunted by low SES.

WM at age 14 also negatively predicted the amount of WM change over adolescence. However, due to some ceiling in our WM tasks at the second time-point (an inherent problem in longitudinal studies), we interpret this result with caution. If this result reflects a true effect, it could represent a catch-up – subjects with lower WM show greater gains in WM during adolescence. However, a previous study of PGS and SES on IQ showed a widening gap between subjects with low and high EduYears-PGS ^45^. At least in part, our results here could alternatively be explained by ceiling effects, which could artificially lead to high performing subjects having less room to improve.

### Limitations and SES subcomponents

As in most studies that use PGS, we were technically restricted to analyzing only subjects of European ancestry, so our results here cannot generalize to other ethnicities. Additionally, our study focused on environmental and genetic predictors of the trait educational attainment. Therefore, our SES measure is “pure” in the sense that we removed some of the genetic variation associated with educational attainment present in the teenagers (and that other studies typically confound). Importantly, however, this controlling does not mean that our SES measure is environmentally pure – there are other potential genetics factors associated with SES, such as genes modulating other traits correlated with SES, as well as parental genes for educational attainment that were not passed down to the teenagers but that helped shaping the environment where they grew up ^46^.

It is also worth noting that the SES composite we used here is a combination of many distinct SES-related components. When splitting this into education, income, and neighborhood components, we found that most of the SES findings in our sample are driven by the level of education of the parents. We were surprised that even after controlling for the best composite of genetic variants currently known for educational attainment (EduYears-PGS from the largest GWAS to date), the education of the parents was still associated with WM and brain structures in our models. Possible environmental factors might be study habits, cognitively stimulating environments or books in a household but might also be diet and stress, all of which are related to higher education. It would be of interest to further study the specific environmental and nontransmitted genetic factors related to cognitive and brain development.

## Conclusions

Here we report, for the first time, distinct associations of EduYears-PGS and SES on cognition, brain structure, and adolescent brain development. These findings imply that behavioral and psychological consequences of SES are likely wide ranging, and less targeted towards a specific cognitive function or behavioral deficit. Importantly, SES has a significant relationship with cognition, even after removing genetic variance. A continued greater insight into the genetics of cognitive development will help inform policy decision to tackle environmental influences.

## Methods

### Study description

IMAGEN is a European multi-site longitudinal genetic and neuroimaging study. All procedures were approved by each of the sites’ (Berlin, Dresden, Dublin, Hamburg, London, Mannheim, Nottingham, Paris) ethics committees. Written informed consent was obtained from the adolescents and parents involved in the study. Our study uses data from the first two neuroimaging waves, at ages fourteen (14.44, SD = 0.38) and nineteen (19.01, SD = 0.72). See Schumann and colleagues^14^ for more information on IMAGEN protocols and inclusion/exclusion criteria.

For subjects to be included in our study, they had to be of European ancestry (due to limitations of the imputations and possible inferences for creating the EduYears-PGS) and have no siblings included in the study (the few sibling pairs were only present due to mistakes in data collection). Importantly, subjects also had to have all of the relevant data available; structural MRI’s at both timepoints, genetics, relevant demographics (e.g., gender and age) and three behavioral WM tasks at the first timepoint. Lastly, genetic and neuroimaging data had to pass their respective quality controls (criterion discussed in depth below). This resulted in a final sample of 551 subjects (321 females).

### Behavioral measures

We estimated working memory (WM) based on three cognitive tasks from the CANTAB battery available in IMAGEN. We combined these into a latent factor that explained around 40% of the common variance. The tasks were: 1) Spatial WM task (SWM), in which participants must search for a token hidden in one of many boxes. The token does not repeat location, and the measure consisted of the number of times participants returned to search a box that had a token. 2) Pattern Recognition Memory task (PRM), in which participants must remember 12 abstract patterns shown in a sequence. The measure consisted of correct choices on a two alternative forced choice task immediately after encoding. 3) Rapid Visual Information Processing task (RVP), in which participants must monitor for a 3-digit target sequence from a stream of 100 digits per minute. The measure used was correct responses.

### Socioeconomic status

The socioeconomic status (SES) score was comprised of the sum of the following variables: Mother’s Education Score, Father’s Education Score, Family Stress Unemployment Score, Financial Difficulties Score, Home Inadequacy Score, Neighborhood Score, Financial Crisis Score, Mother Employed Score, Father Employed Score (see SI Appendix, Table S4). To further disentangle SES-related variation, post hoc we divided our measure into three summed sub-components: Parental Education (comprised of mother’s and father’s education score), Neighborhood related factors (comprised of neighborhood score and home inadequacy score) and Income-related variables (comprised of financial difficulties score, financial crisis score and family unemployment stress).

### EduYears Polygenic Score

All participants in IMAGEN had DNA extracted from blood samples and were genotyped with the Illumina Human610-Quad Beadchip or the Illumina Human660-Quad Beadchip. A PCA approach was used to identify and exclude individuals with non-European ancestry. The quality control procedures excluded SNPs with call rates >95%, minor allele frequencies less than 5%, and SNPs that did not pass an exact test of Hardy-Weinberg equilibrium at P < 5×10^−4^. After quality control, around 480,000 SNPs were then used for imputations via a reference file created by the ENIGMA2 Genetics Support Team. Haplotype phasing and imputation was performed using, respectively, Mach1 and Minimac codes from the MaCH software suit ^47^, as specified in the ENIGMA2 protocol (http://enigma.ini.usc.edu/wpcontent/uploads/2012/07/ENIGMA2_1KGP_cookbook_v3.pdf).

We then used this genotype data to estimate EduYears-PGS in each participant based on the effect sizes of thousands of SNPs discovered by the most recent GWAS on educational attainment ^13^. We first obtained the summary statistics of the GWAS from the Social Science Genetic Association Consortium (https://www.thessgac.org/data). We decided which significance threshold to use in our sample by performing high resolution scoring in PRSice-2 ^48^ based on the phenotype of WM (from the latent factor of our three cognitive tasks). The threshold that optimally explains the variance in WM resulted in an EduYears PGS containing 5709 SNPs. To guard against overfitting, we performed 1 million permutations and obtained a significant empirical p-value of our estimate (empirical p = 0.009). EduYears-PGS in our sample was standardized to have a mean of zero and a standard deviation of one for the population. All steps for creating the EduYears-PGS were performed using the R package PRSice-2 ^48^ and PLINK version 1.90 ^49^.

### Structural Imaging

#### Image Acquisition and Preprocessing

Structural imaging data were acquired using numerous 3T MRI scanners (Philips Medical Systems Achieva, Bruker, Siemens TrioTim, Siemens Verio, Bruker/GE Medical Systems Signa Excite, GE Medical Systems Signa HDx) with a T1-weighted gradient echo sequence (isotropic 1.1mm) based on the ADNI protocol (http://adni.loni.usc.edu/methods/documents/mri-protocols) ^50^. Freesurfer (v6.0.0; http://surfer.nmr.mgh.harvard.edu/) was used for image preprocessing and SA/CT estimation, previously reported in depth ^51^. Specifically, the longitudinal pipeline was used as it is optimized for longitudinal data by registering the differing timepoints to a median image thereby reducing within-subject variability and avoiding registration bias ^52^. All processing was run using the high-performance computing (Bianca cluster) resources provided by SNIC through Uppsala’s Multidisciplinary Center for Advanced Computational Science (UPPMAX) under Project sens2018615 using gnuparallel ^53^. Since IMAGEN is a multi-center study we used ComBat at the vertex-wise level to remove unwanted site-based variability ^54^. Variables of interest (baseline age, EduYears-PGS, SES, scan interval, gender) also showed site specific variability, therefore we entered these in ComBat to retain these true site-specific differences. Mean CT and SA were calculated by averaging all of the vertices. For vertex-wise analysis we smoothed the data with a gaussian smoothing kernel of 10mm FWHM.

#### Quality Control

The IMAGEN team provided anatomical quality control for the second time-point and partially for the first time-point. Following longitudinal preprocessing, two raters graded the Desikan–Killiany parcellated atlas and the white matter/pial boundary overlaid on the T1 (norm.mgz) in a coronal view separately for both the final longitudinal timepoints and the base (median) image on a 3-point scale (pass, doubtful, fail). Any scan that was marked ‘doubtful’ was reviewed by the other rater, and a consensus decision to include (pass) or exclude (fail) was made. A large number of scans were determined to have skull strip errors (via the pial boundary). As previous research on cortical thickness has shown that quality control can impact the conclusions drawn ^41^, we reran recon-all on a subset of subjects showing skull strip errors with ‘gcut’. If the error persisted in either timepoint the subject was excluded. From 1963 subjects, 1168 had structural scans at both timepoints, from these 748 passed quality control (pass rate 64%). In a similar fashion to previous research with large sample sizes quality control was not entirely overlapping between the two raters ^55^. In the overlap sample of 101 subjects we found a high inter-rater agreement (kappa = 0.88).

### Statistical analysis

#### Surface area, Cortical thickness, Hippocampus & Amygdala

We choose to use a bivariate latent change score (bLCS) model as it allowed us to examine the development of neural and behavioral measures simultaneously without the constraints of measurement error ^56,57^. A latent change score model can be conceptualized as a reparameterization of a paired t-test and has recently been highlighted for its usefulness in teasing apart the complex processes involved in longitudinal developmental research ^58,59^. bLCS models were estimated for average cortical thickness and total surface area. For scaling purposes, we defined total surface area as the sum of all vertices divided by the amount in standard space. For model estimation we used full information maximum likelihood and a robust maximum likelihood estimator with a Yuan-Bentler scaled test statistic from the R (v. 3.6.0) package Lavaan (v. 0.6-3) ^60,61^. Missing follow-up behavioral data was imputed under the assumption of missing at random (see SI Appendix, Fig. S3) ^62^. We assessed model fit using the comparative fit index (CFI; fit > 0.95) and the root mean square error of approximation (RMSEA; fit < .08) ^63^. The subjects’ age and gender were regressed from all observed behavioral and neural measures in both timepoints before model fitting. For the hippocampus and amygdala analyses total intercranial volume was also corrected for. All models have strict measurement invariance; intercepts, loadings and error variance were constrained to be equal across time. Prior to fitting a bLCS model we assessed fit on respective measurement models. If estimates present Heywood cases (negative error variances) we left them unconstrained as long as the null hypothesis could not be rejected with the use of confidence intervals ^64^. Any model presented will have a positive upper bound confidence interval; we chose this approach since constraining variance can lead to unintended consequences ^64,65^. Each SES sub-component was individually substituted for the SES composite in all models, p-values were FDR corrected for the novel tests (see SI Appendix, Table S5).

#### Vertex-wise exploratory analysis

An inherent goal in vertex-wise exploratory analysis is anatomical localization therefore we corrected for global values (i.e., total surface area and average cortical thickness). Correcting for global values in the analysis at age 14 allowed us to detect regional areas that are unrelated to the overall global effects (see discussion for further elaboration). Linear models were fit for each vertex in Freesurfer using Monte Carlo based cluster-wise correction with a cluster-forming threshold of .001, a cluster-wise alpha of .05 and Bonferroni correction for making two independent tests for the two hemispheres ^66,67^.

We used two linear models to probe regional specificity of SES and EduYears-PGS, one for subjects at 14 and another on the vertex-wise subtracted images (age 19 subtracted from age 14). Both models were fit for SA and CT separately, resulting in a total of 4 analyses. In the model at age 14 vertices were predicted by the EduYears-PGS and SES while being controlled for mean value of the modality analyzed, gender, age at the first scan and gcut. Gcut is a dummy variable coding for subjects who needed stricter skull strip processing to pass quality control. Subtracted vertices were predicted by SES and EduYears-PGS while being controlled for gender, between scan interval and gcut.

#### Data availability

It was not part of the written consent of the participants for the data to be publicly shared. Researchers may access the dataset through a request to the IMAGEN consortium: https://imagen-europe.com/resources/imagen-project-proposal/

## Acknowledgments

This work received support from the following sources:

Vetenskapsradet (Swedish research council; 2015-02850), Wenner-gren foundation (UPD2018-0295), National Institute of Health: (Meaningful Data Compression and Reduction of High-Throughput Sequencing Data”) (1 U01 CA198952-01), The European Union-funded FP6 Integrated Project IMAGEN (Reinforcement-related behaviour in normal brain function and psychopathology) (LSHM-CT-2007-037286), the Horizon 2020 funded ERC Advanced Grant ‘STRATIFY’ (Brain network based stratification of reinforcement-related disorders) (695313), ERANID (Understanding the Interplay between Cultural, Biological and Subjective Factors in Drug Use Pathways) (PR-ST-0416-10004), BRIDGET (JPND: BRain Imaging, cognition Dementia and next generation GEnomics) (MR/N027558/1), Human Brain Project (HBP SGA 2, 785907), the FP7 project MATRICS (603016), the Medical Research Council Grant ‘c-VEDA’ (Consortium on Vulnerability to Externalizing Disorders and Addictions) (MR/N000390/1), the National Institute for Health Research (NIHR) Biomedical Research Centre at South London and Maudsley NHS Foundation Trust and King’s College London, the Bundesministeriumfür Bildung und Forschung (BMBF grants 01GS08152; 01EV0711; Forschungsnetz AERIAL 01EE1406A, 01EE1406B), the Deutsche Forschungsgemeinschaft (DFG grants SM 80/7-2, SFB 940, TRR 265, NE 1383/14-1), the Medical Research Foundation and Medical Research Council (grants MR/R00465X/1 and MR/S020306/1), the National Institutes of Health (NIH) funded ENIGMA (grants 5U54EB020403-05 and 1R56AG058854-01). Further support was provided by grants from: - ANR (project AF12-NEUR0008-01 - WM2NA, ANR-12-SAMA-0004), the Eranet Neuron (ANR-18-NEUR00002-01), the Fondation de France (00081242), the Fondation pour la Recherche Médicale (DPA20140629802), the Mission Interministérielle de Lutte-contre-les-Drogues-et-les-Conduites-Addictives (MILDECA), the Assistance-Publique-Hôpitaux-de-Paris and INSERM (interface grant), Paris Sud University IDEX 2012, the fondation de l’Avenir (grant AP-RM-17-013), the Fédération pour la Recherche sur le Cerveau; the National Institutes of Health, Science Foundation Ireland (16/ERCD/3797), U.S.A. (Axon, Testosterone and Mental Health during Adolescence; RO1 MH085772-01A1), and by NIH Consortium grant U54 EB020403, supported by a cross-NIH alliance that funds Big Data to Knowledge Centres of Excellence.

